# Enzymatic Function of an Intrinsically Disordered Protein

**DOI:** 10.1101/2025.09.02.673730

**Authors:** Tatiana Lyalina, Lia Mara Gomes Paim, Susanne Bechstedt

## Abstract

Intrinsically disordered proteins (IDPs) challenge the traditional structure-function paradigm by lacking a stable three-dimensional structure ^1^. While their roles as dynamic effectors, scaffolds, and molecular switches are well-established, it has been widely accepted that enzymatic activity requires a stably folded catalytic center ^2^. Here, we challenge this dogma by demonstrating that a 284-amino acid intrinsically disordered domain of the cytoskeleton-associated protein 2 (CKAP2) is sufficient to catalyze both microtubule polymerization and depolymerization. CKAP2 promotes tubulin incorporation without high-affinity tubulin binding, suggesting a transition-state-based catalytic mechanism distinct from known microtubule polymerases. These findings establish, for the first time, that an intrinsically disordered domain can function as a bona fide enzyme, expanding our understanding of the functional repertoire of disordered proteins and their roles in cellular processes.

## Main

Intrinsically disordered proteins (IDPs) lack a stable three-dimensional structure under physiological conditions, yet they carry out a wide range of essential biological functions ^1^. In the human proteome, nearly one-third of amino acids are found within intrinsically disordered regions (IDRs) ^3^. These flexible domains act as scaffolds, molecular switches, and dynamic effectors in signalling pathways, often through transient, low-affinity interactions ^1^. More recently, IDRs have also been implicated in driving biomolecular condensation via multivalent interactions governed by a “molecular grammar” ^4–7^. Despite this functional diversity, no intrinsically disordered domain has been shown to exhibit enzymatic function, an activity long thought to require the presence of a stably folded catalytic center ^2^.

Microtubules, dynamic polymers composed of α/β-tubulin heterodimers, are crucial for fundamental cellular processes, including mitosis, intracellular transport, and cell polarity ^8–10^. Their rapid turnover, occurring on the order of seconds in dividing cells and minutes in neurons ^11–13^, is tightly regulated by microtubule-associated proteins (MAPs) ^14,15^. Among these, the conserved chTOG/XMAP215/Stu2 family acts as microtubule polymerases, using conserved α-helical repeats, known as TOG domains, which bind free tubulin with high affinity ^16–19^. This activity is thought to involve a tethered or “antenna-like” mechanism that concentrates tubulin subunits at the microtubule plus end to deliver tubulin dimers to growing microtubule ends ^16–19^. Similarly, we have previously reported that the Cytoskeleton-associated protein 2 (CKAP2) also promotes microtubule growth *in vitro* and in cells ^20,21^. However, its mechanism of action and structural organization remain unknown, as the protein largely lacks defined structural domains.

Here, we demonstrate that a central intrinsically disordered domain of CKAP2 is sufficient to catalyze both microtubule polymerization and depolymerization. These findings establish, for the first time, that a disordered domain can function as an enzyme.

## Results

We have previously shown that approximately 60% of CKAP2 is disordered ^21^. To investigate its structural organization, we used MetaPredict ^22^ to identify disordered and ordered domains within the full-length protein (Fig. 1a). We also applied AlphaFold3 to predict the structure of CKAP2 and model its interaction with two tubulin dimers (Fig. 1b) ^23^. These analyses identified four distinct domains within CKAP2 (Fig. 1b,c and Supplementary Video 1). The AlphaFold3 model predicts that the N-terminal α-helix, the Disordered domain, and the central α-helical bundle interact with tubulin. In this model, the structured C-terminal domain does not contact tubulin, while among the three potential tubulin-binding domains, only the Disordered domain is predicted to bind to sites accessible within the fully formed microtubule lattice. In contrast, the predicted contact sites for the N-terminal α-helix and the central α-helical bundle domain are at least partially buried within polymerized microtubules, suggesting that these domains may only interact with unpolymerized tubulin dimers.

**Fig. 1:**
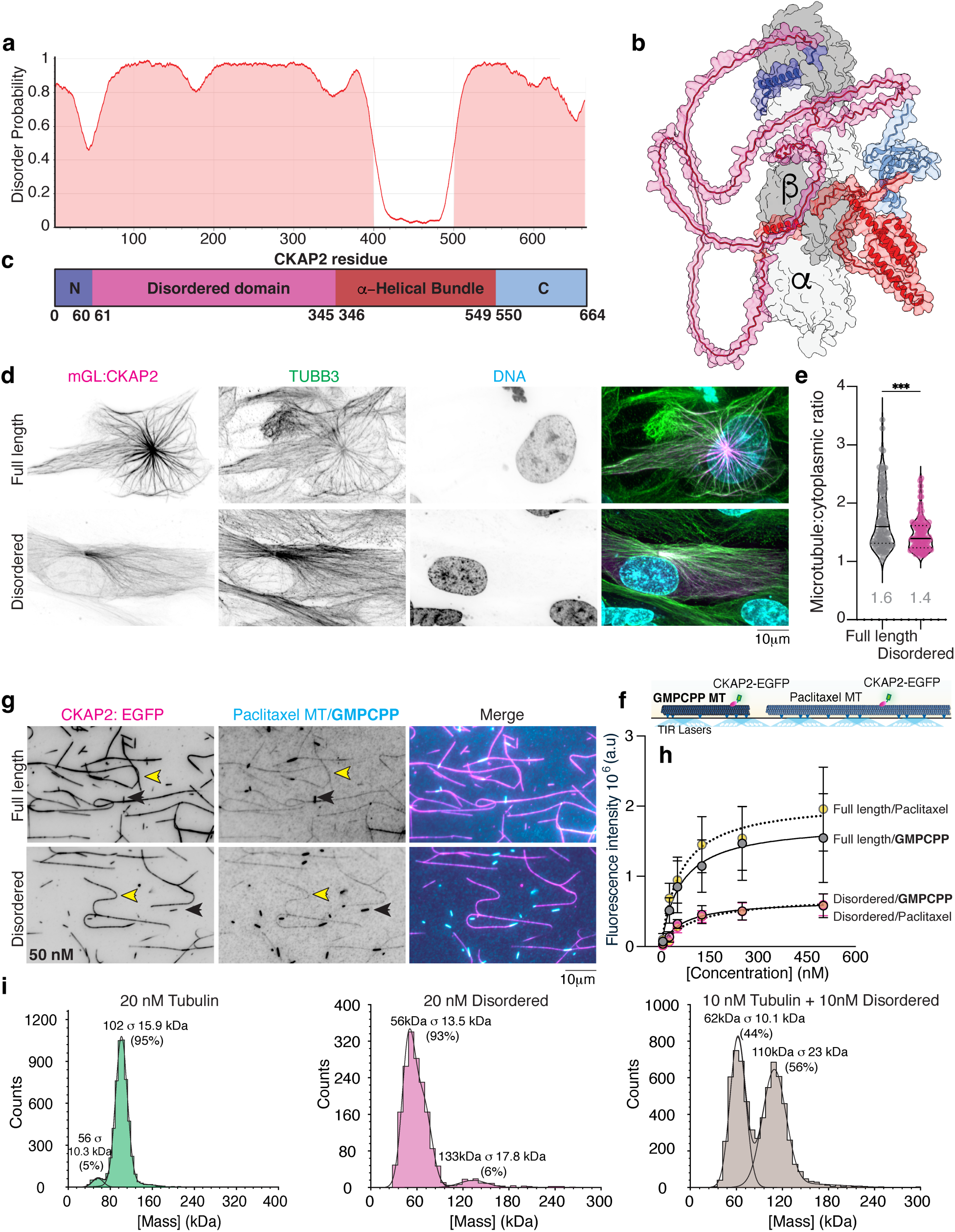
The Disordered domain in CKAP2 is responsible for microtubule binding. **a,** MetaPredict disorder prediction for CKAP2. **b,** AlphaFold3 Model for CKAP2 binding to two tubulins. **c,** Schematic of domains identified through AlphaFold3: the N-terminal α-helix (in dark blue), the Disordered domain (in pink), the central α-helical bundle (in red), and the C-terminus (in light blue). **d,** Representative images for full-length CKAP2 and the Disordered domain binding to microtubules in cells. **e,** Quantification for microtubule binding in cells. Full length n = 110 microtubules from 22 cells, disordered n = 115 microtubules from 23 cells. *P* value is indicated *** = 0.0002 (Mann-Whitney test). Values depicted in the chart refer to the median. **f,** Schematic of *in vitro* microtubule binding assay. **g,** Representative images for full-length CKAP2 and the Disordered domain binding to paclitaxel and GMPCPP-stabilized microtubules at 50 nM. Yellow arrows highlight dim paclitaxel microtubules; black arrows indicate bright GMPCPP microtubules. **h,** Binding curves for data in g, fitting to a hyperbola. N microtubules vary from 32 to 50. Circles represent the mean of each concentration for full-length CKAP2 and the Disordered domain binding to paclitaxel and GMPCPP-stabilized microtubules. **i,** Mass Photometry data for tubulin and the Disordered domain, separate and together.

### AlphaFold model Validation

To validate the AlphaFold3 model, we expressed the four CKAP2 domains fused to GFP in human RPE-1 cells to probe their interactions with cellular microtubules (Fig. 1d). Consistent with our predictions, only the Disordered domain exhibited a high-affinity interaction with microtubules in cells (Fig. 1d,e and Extended Data Fig. 1h,i). The expression of the Disordered domain resulted in a slightly lower cytoplasmic-to-microtubule ratio compared to the full-length protein, suggesting a lower affinity for microtubules (Fig. 1e). Its expression also led to the accumulation of stable microtubules visible as bundles, similar to those of full-length CKAP2, a characteristic of its polymerizing activity or role in stabilization (Extended Data Fig. 1j,k).

Additionally, we expressed the four domains fused to GFP in *E. coli* (Extended Data Fig. 1a) and visualized the interaction of these purified proteins with reconstituted GMPCPP- and paclitaxel-stabilized microtubules in an *in vitro* microtubule binding assay (Fig. 1f,g). This assay confirmed the interaction of the Disordered domain with microtubules and that its affinity to microtubules is significantly reduced compared to full-length CKAP2 (Fig. 1h). The other three domains showed no interaction with microtubules in cells (Extended Data Fig. 1h-k). The N-terminal α-helix and the central α-helical bundle also show no significant interaction with *in vitro* polymerized microtubules (Extended Data Figures 1c,d). Unlike the AlphaFold3 predictions, the C-terminal domain exhibited a low-affinity interaction with microtubules *in vitro* (Extended Data Fig. 1c,d), which we suspected could be due to the interaction of positively charged patches in this domain with the negatively charged tubulin tails (Extended Data Fig. 1e). Indeed, we found that the C-terminal domain ceases to bind microtubules when the tubulin C-terminal tails are removed through limited digestion with the protease subtilisin (Extended Data Figures 1f,g).

Next, we examined the interaction of CKAP2 domains with free tubulin using mass photometry, a label-free single-molecule technique that measures the molecular mass of proteins in solution based on light scattering and detects sub-micromolar affinities ^24^. In this assay, none of the CKAP2 domains showed any interaction with free tubulin, unlike the positive control, the TOG-domain polymerase Stu2 (Fig. 1i and Extended Data Fig. 1b). This finding is consistent with previous results indicating that there is no interaction of the full-length CKAP2 in size-exclusion chromatography ^21^. Together, these results rule out a high-affinity interaction between CKAP2 and free tubulin, suggesting that its mechanism of action differs from that of TOG-domain polymerases, which bind tubulin with an affinity of 10 nM ^16^.

### The CKAP2 IDR contains all core functionality

To confirm that the Disordered domain (amino acids 61-345) is indeed intrinsically disordered, we performed circular dichroism (CD) spectroscopy (Fig. 2a). Analysis of the CD spectra indicates a secondary structure content of approximately 5% α-helix and 2% β-sheet, a value that falls within the error margin of the fit and is not statistically distinguishable from zero. This data confirms that this domain lacks any meaningful secondary and tertiary structure elements and is intrinsically disordered in solution.

**Fig. 2:**
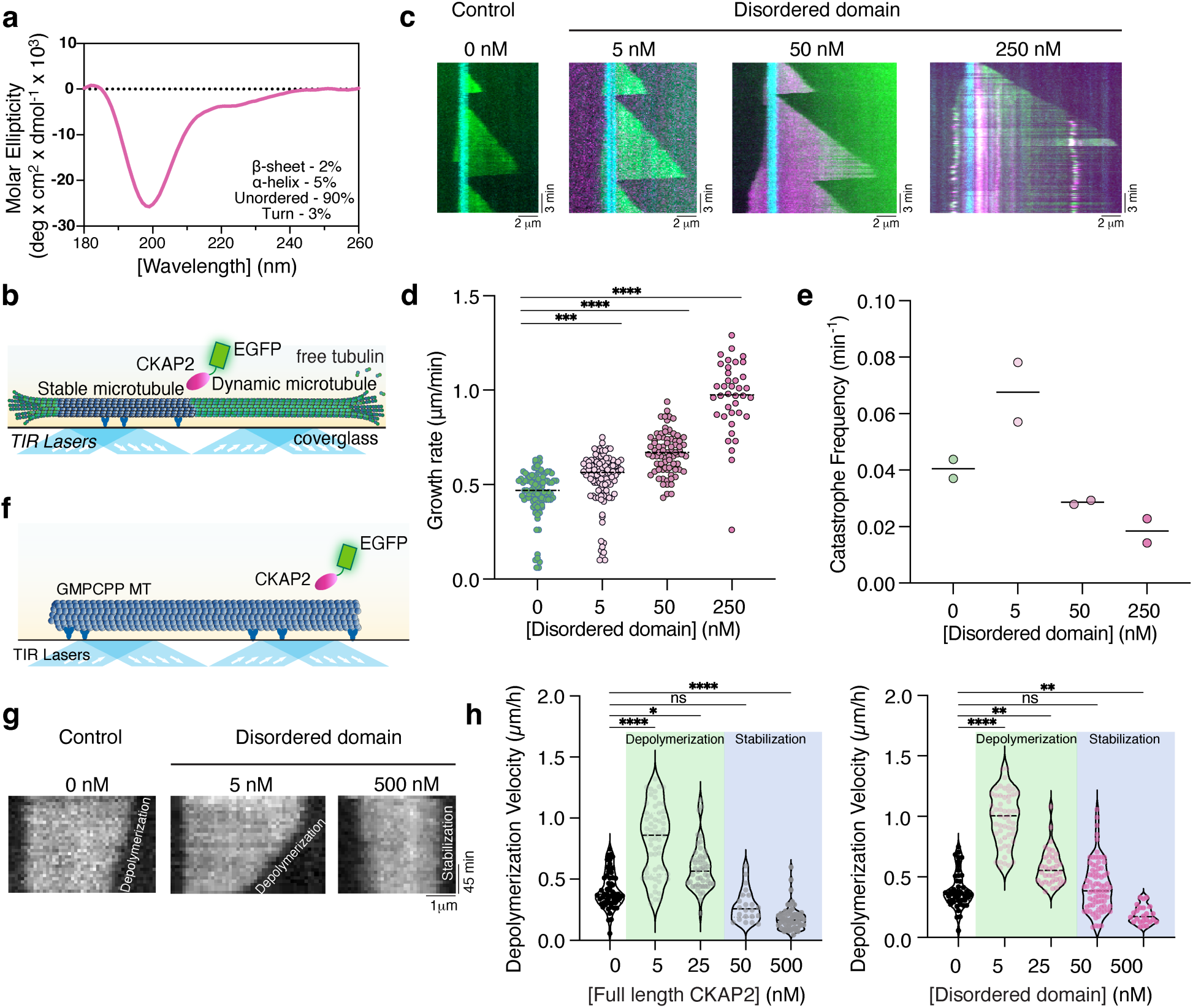
The Disordered domain retains enzymatic function. **a,** Circular dichroism of the Disordered domain of CKAP2. **b,** Schematic of the dynamic microtubule assay. **c,** Representative kymographs of microtubule growth in the absence (0 nM) and presence of 5, 50, and 250 nM of the Disordered domain of CKAP2. **d,** Quantification of microtubule growth rates for data in c. N = 97, 102, 78, 38 growth events for 0, 5, 50 and 250 nM, respectively. The dashed black bar denotes the median 0.47, 0.57, 0.67, 0.98 for 0, 5, 50 and 250 nM from at least 2 independent replicates. *P* values are indicated *** = 0.0003, **** < 0.0001, **** < 0.0001 (Kruskal-Wallis test). **e,** Quantification of microtubule catastrophe frequency for data in c. *P* values are indicated ** = 0.0038, ns > 0.9999, ns = 0.1729 (Kruskal-Wallis test). **f,** Schematic for the microtubule depolymerization assay. **g,** Representative kymographs of GMPCPP-stabilized microtubule depolymerization in the presence of 0, 5 and 500 nM of the Disordered domain. **h,** Quantification of GMPCPP microtubule depolymerization for full-length CKAP2 and the Disordered domain. N analyzed microtubules vary from 22 to 79. At least three independent experiments were done. *P* values are indicated for the Full length **** < 0.0001, * = 0.0117, ns = 0.2035, **** < 0.0001 and for the Disordered region ****< 0.0001, ** = 0.0039, ns > 0.9999, ** = 0.0030 (Kruskal-Wallis test). The dashed black bar indicates the median.

To determine if, in addition to microtubule binding, the Disordered domain alone retains any functional activity of CKAP2 on microtubules, we used an *in vitro* reconstitution assay for microtubule dynamics (Fig. 2b). We observed the effect of varying concentrations of the Disordered domain on microtubule growth and shrinkage (Fig. 2c and Supplementary Video 2). Remarkably, the Disordered domain alone was sufficient to promote microtubule growth at concentrations as low as 5 nM, and doubled the polymerization rate compared to controls at 250 nM (Fig. 2d). The Disordered domain also significantly stabilized microtubules at concentrations above 50 nM, reducing the catastrophe frequency by 50% at 250 nM (Fig. 2e). These findings demonstrate that the Disordered domain contains the core functionalities of CKAP2.

### The CKAP2 IDR is an Enzyme

We previously observed that CKAP2 promotes microtubule growth, but the mechanistic basis for this activity was not fully elucidated ^21^. Considering that TOG domain polymerases serve as established models for microtubule growth-accelerating enzymes ^16,17^, we investigated whether CKAP2 exhibits similar enzymatic characteristics. A defining property of enzymes is their ability to catalyze the reverse reaction in the absence of a substrate ^6,17,19^. To test whether CKAP2 exhibits this hallmark of enzymatic activity, we examined its ability to promote microtubule depolymerization in the absence of free tubulin. Using microtubules stabilized with the slowly hydrolyzable GTP analog GMPCPP, we tracked depolymerization over 2.5 hours (Fig. 2f,g and Supplementary Video 3). We observed that both full-length CKAP2 and the Disordered domain accelerated depolymerization rates approximately twofold at 5 nM and to a lesser extent at 25 nM (Fig. 2h). At higher concentrations (500 nM), both full-length CKAP2 and the Disordered domain suppressed depolymerization, consistent with our dynamic assay data, which showed reduced catastrophe frequency and shrinkage rate at 250 nM (Fig. 2e and Extended Data Fig. 2a). This experiment provides definitive evidence that both full-length CKAP2 and the Disordered domain catalyze the reverse reaction of tubulin polymerization in the absence of substrate and therefore function as enzymes. At higher lattice occupancy, this enzymatic activity is masked by the lattice-stabilizing functionality of CKAP2.

## Discussion

Our results demonstrate that CKAP2 functions as a bona fide enzyme, with its activity residing entirely within an intrinsically disordered domain. This discovery challenges the long-standing paradigm that enzymatic activity requires a stably folded catalytic center.

We show that CKAP2 is an effective polymerase and depolymerase at low nanomolar concentrations. Higher concentrations that increase lattice occupancy additionally confer microtubule stabilization and protect against depolymerization. These dual effects are recapitulated by the isolated disordered domain, indicating that it harbours the complete functional repertoire of CKAP2. While the disordered domain is sufficient for activity, additional positively charged patches in the full-length protein enhance microtubule affinity, a common feature among microtubule-associated proteins (MAPs) and intrinsically disordered proteins (IDPs) ^25,26^. Notably, lysine residues constitute approximately 10% of CKAP2’s sequence, twice the average found in typical eukaryotic proteins ^27^. Positive patches are found particularly in the N and C-terminal domains, explaining the low residual affinity of these domains for microtubule lattices (Extended Data Fig. 1e). The highly polybasic nature of CKAP2 is consistent with its strong interaction with the negatively charged microtubule C-terminal tails.

Our findings demonstrate that the Disordered domain of CKAP2 lacks defined secondary structural elements and does not adopt a folded three-dimensional conformation, as supported by structural analysis (Fig. 2a). While some intrinsically disordered proteins (IDPs) fold upon binding their targets ^28,29^, predictive modelling suggests that this is not the case for CKAP2.

Notably, AlphaFold3 does not predict the formation of any structured elements within the CKAP2 binding region ^23^. Moreover, AlphaFold-Multimer, which achieves approximately 76% accuracy in modelling IDR interactions ^30,31^, also fails to predict stable structure formation in the CKAP2-tubulin complex. Together, these observations strongly suggest that CKAP2 remains disordered and does not acquire secondary or tertiary structure upon binding to microtubules or tubulin.

Overall, our findings reveal that CKAP2 promotes microtubule polymerization through a mechanism fundamentally distinct from that of canonical TOG domain-containing polymerases such as those in the chTOG/XMAP215/Stu2 family. First, we observed no high-affinity interaction with free tubulin, a key feature of TOG domains ^16^. This indicates that CKAP2 does not function as a “tubulin antenna” like TOG domain polymerases, and reagent concentration is not part of the mechanism by which CKAP2 operates. Instead, we propose that CKAP2 targets the microtubule end, where it stabilizes the transition state of an incoming tubulin dimer. By lowering the activation energy required for both the addition and dissociation of tubulin subunits, CKAP2 catalyzes both polymerization and depolymerization reactions. Structurally, this may involve aiding the transition of tubulin from a curved to a straight conformation or the formation of longitudinal and lateral inter-tubulin dimer bonds ^32,33^. Indeed, it has recently been suggested that the transition state could be modified by regulatory factors of microtubule growth ^33^. At higher concentrations of CKAP2, higher lattice occupancy of CKAP2 stabilizes the lattice against catastrophe and shrinkage in growth- and depolymerization assays (Fig. 2g,h and Extended Data Fig. 2a).

We propose that the 21-amino-acid microtubule-binding segment within CKAP2 functions as a short linear motif (SLiM). SLiMs are well-established effector regions within intrinsically disordered regions (IDRs) ^34–36^. The CKAP2 SLiM shows strong evolutionary conservation (Extended Data Fig. 3a,b), highlighting its functional importance. The inherent flexibility of disordered regions provides an advantage over folded domains by lacking a tightly-bound structured substrate binding state ^1,37^. For a microtubule growth factor, this flexibility allows rapid molecular recognition of the transition state of the incoming tubulin dimer and efficient unbinding, facilitating the release of dimers once incorporated and freeing CKAP2 for the next addition cycle (Fig. 3a). Consistent with this mechanism, CKAP2 surpasses traditional TOG domain-based polymerases in accelerating microtubule growth. While XMAP215 increases the tubulin on-rate (k_a_) by up to 5-fold, CKAP2 achieves a striking 50-fold increase ^17,21^.

**Fig. 3:**
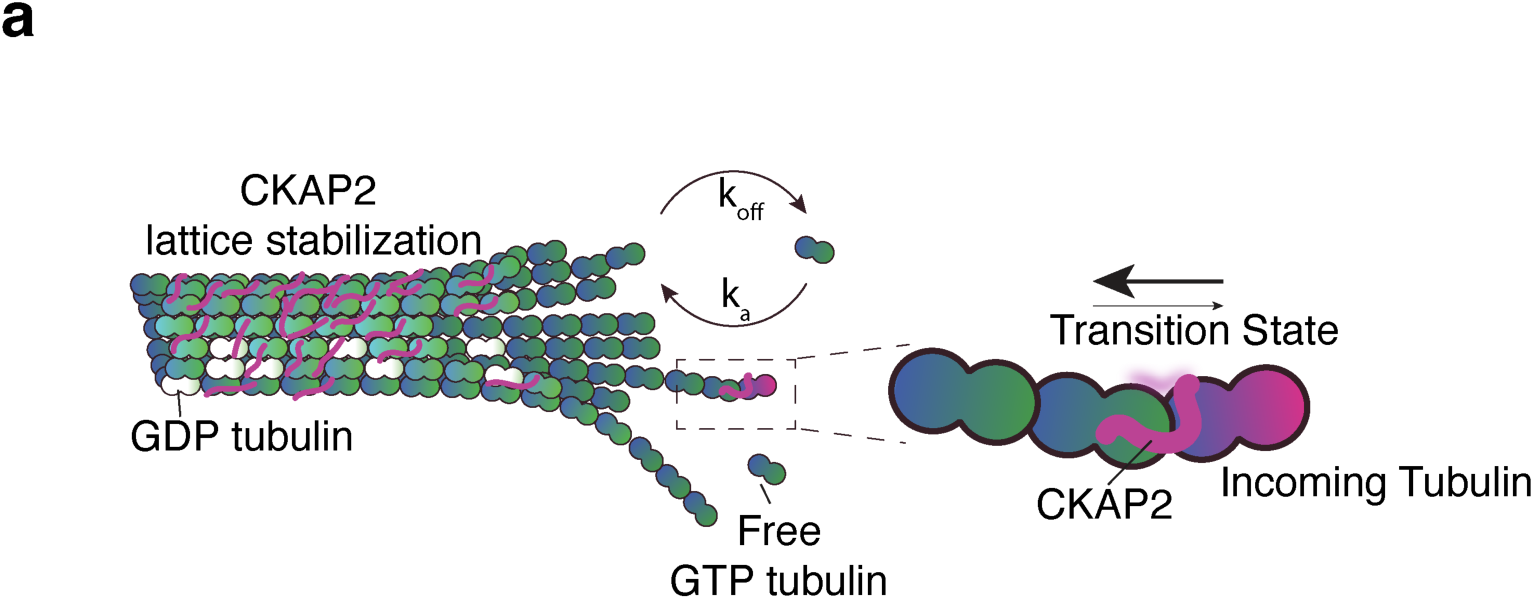
The disordered domain of CKAP2 functions as a disordered enzyme. **a,** Model for CKAP2’s mechanism in microtubule polymerization and stabilization.

SLiMs also provide a mechanism for the precise regulation of protein-protein interactions through post-translational modifications (PTMs) ^38^. In CKAP2, phosphorylation of the last conserved serine residue within the binding motif has been identified as a prominent modification site ^39^. The introduction of a negative charge through phosphorylation at this position will substantially weaken microtubule binding and polymerization activity. Given that CKAP2 is a potent microtubule assembly factor and reaches cellular concentrations in the hundreds of nanomolar range during mitosis ^40^, tight temporal and spatial control of its activity is likely essential for proper spindle formation and genome stability ^20^.

In summary, our study offers the first direct evidence of enzymatic activity within an intrinsically disordered protein domain, marking a paradigm shift that broadens our understanding of the functional diversity of IDPs. Our work also provides a new mechanism for modulating cytoskeletal filament dynamics. This work opens new avenues for exploring the catalytic potential of other disordered regions and their roles in fundamental biological processes and diseases.

## Methods

### Structure Prediction

Disorder prediction for full-length CKAP2 (UniProt accession number Q3V1H1) was performed using Metapredict ^22^. For three-dimensional structure prediction and protein-protein interaction modeling, AlphaFold3 ^23^ was utilized. In addition to CKAP2, two copies each of ubiquitous human α-tubulin 1A (UniProt accession number Q71U36) and β-tubulin 1 (UniProt accession number Q9H4B7) were included in the AlphaFold3 predictions to model their interactions. Molecular graphics and analyses were performed with the UCSF Chimera package (ChimeraX 1.8) ^41^.

Amino acid sequences for CKAP2 from different species were obtained from UniProt, aligned with ClustalW, and then visualized using MacVector 17.0.10.

### Cloning

Full-length CKAP2 was obtained as described previously ^21^. CKAP2 domains were amplified from the full-length plasmid using Pfux7 polymerase ^42^ and inserted into either a pHAT vector containing an N-terminal 6-His-tag, followed by the protein of interest, EGFP, and a Strep-tag II for protein expression ^43,44^ or a pTWIST EF1alpha-Puro (Twist Bioscience) for mammalian expression. The primers were ordered from IDT Technologies and are listed in Supplementary Table 1. The disordered domain was also cloned into a pHAT vector without EGFP. *Saccharomyces cerevisiae* Stu2 Δcc aa 1-648 was amplified from yeast genomic DNA (a gift from Daici Chen/Jackie Vogel, McGill University). Plasmid DNA was transformed into NEB Turbo competent *E. coli*, and minipreps were prepared using Presto™ Mini Plasmid Kit (PDH300) according to the manufacturer’s instructions (Geneaid, Taiwan). Correct inserts were confirmed by Sanger sequencing.

### Circular Dichroism

Circular dichroism (CD) spectra were collected for assessing the secondary structural content of the purified unlabelled disordered domain of CKAP2 (3.12 μM) in Tris buffer (100 mM Tris-HCl, 150 mM NaCl, 1 mM EDTA, 2.5 mM Desthiobiotin, pH 7.8) at room temperature using a Chirascan CD Spectrometer (Applied Photophysics Ltd., Leatherhead, Surrey. U.K.).

Twelve replicate circular dichroism scans were collected with 1 nm increments and 0.5-second integration times using a 0.1 mm pathlength sample cuvette, spanning the wavelength range from 180 nm to 260 nm. Raw spectra were corrected with reference spectra for the corresponding buffer and temperature. These corrected spectra were then processed with 3-point arithmetic smoothing using the Chirascan ProViewer software (version 4.7.0.194). Spectral ranges for each buffer condition were limited to those wavelengths where total sample absorbance did not exceed 1.0 as measured using the Chirascan instrument. Preliminary data processing and reference subtraction were performed with ProViewer software. Secondary structural deconvolution of reference-corrected spectra for the disordered domain of CKAP2 was performed using the CONTINLL algorithm and SPD48 base spectra library within OLIS SpectralWorks software (On-line Instrument Systems, Bogart, Georgia). Two measurements were performed to demonstrate reproducibility.

### Mass photometry measurements of CKAP2 fragments, Stu2 Δcc and tubulin

Mass photometry measurements were performed using a Refeyn TwoMP mass photometer. A calibration curve was obtained using either sweet potato β-amylase (Sigma-Aldrich) or Bovine Serum Albumin (Sigma-Aldrich) diluted in BRB80 buffer (80 mM PIPES-KOH, pH 6.9 (PIPES Sigma BioPerformance, P1851), 1 mM EGTA, 1 mM MgCl_2_) to correlate ratiometric contrast to molecular mass. For measurements, 200 nM of CKAP2 fragments, Stu2 Δcc, and tubulin in BRB80 were kept on ice prior to final dilution. At the moment of MP measurements, 2 µL of either individual proteins (tubulin, CKAP2 fragments, or Stu2 Δcc) or mixtures (CKAP2 fragments + tubulin, Stu2 Δcc + tubulin) were pipetted into 18 µL of BRB80 at room temperature (23°C) on a glass coverslip MassGlass UC slide (Refeyn) to a final concentration of 10 or 20 nM in the mixture. All measurements were done at least in two replicates. Mass histograms were binned, fitted with Gaussian curves, and analyzed in DiscoverMP software (v2024 R1, Refeyn).

### TIRF Microscopy

Images were acquired with a customized Zeiss Axio Observer 7 equipped with a Laser TIRF III and 405/488/561/638 nm lasers, Alpha Plan-Apo 100 ×/1.46 Oil DIC M27, and Objective Heater 25.5/33 S1. Images were recorded using a Prime 95B CMOS camera (Photometrics) with a pixel size of 107 nm, and Zeiss Zen Blue 2.5 was employed for image acquisition.

### Protein expression and purification

The CKAP2 fragments and Stu2 Δcc were expressed in BL21(DE3) *E. coli* grown to OD_600_ = 0.4-0.6 at 37°C and were induced using 0.5 mM Isopropyl ß-D-1-thiogalactopyranoside (IPTG) at 18°C overnight. Bacterial cells were harvested by centrifugation at 2500 g for 10 minutes, resuspended in Buffer A (50 mM NaH_2_PO4, 300 mM NaCl, 0 mM imidazole, pH 7.8), and then lysed in an EmulsiFlex-C5 (Avestin Inc, Canada). Lysates were cleared by centrifugation at 30,000 g for 1 hour and then loaded onto His60 Ni SuperFlow Resin (Takara Bio, USA) in gravity flow columns. The resin was eluted with Buffer B (50 mM NaH_2_PO4, 300 mM NaCl, 200 mM imidazole, pH 7.8). The eluate was loaded onto Strep-Tactin Sepharose resin (IBA Lifesciences, Germany), washed with 5 column volumes of Buffer W (100 mM Tris-HCl, 150 mM NaCl, 1 mM EDTA, pH 7.8) and eluted with Buffer W supplemented with 2.5 mM Desthiobiotin. Protein concentration was determined by absorbance at 280 or 488 nm with a DS-11 FX spectrophotometer (DeNovix Inc, USA). SDS-PAGE was used to determine the purity of the proteins. All the proteins mentioned above were purified at least three times independently to show reproducibility, except for Stu2 Δcc.

Tubulin was purified from bovine brains as previously described ^45^, with the modification of using Fractogel EMD SO3- (M) resin (Millipore-Sigma) instead of phosphocellulose. Tubulin was labelled using CF640R-NHS-Ester (Biotium, USA) and tetramethylrhodamine (TAMRA, Invitrogen Inc, Canada), as described ^46^. An additional cycle of polymerization/ depolymerization was performed before use. Protein concentrations were determined using a DS-11 FX spectrophotometer.

### Channel preparation

Cover glass (22 × 22 and 18 × 18 mm) was cleaned in acetone, followed by sonication in 50% methanol in 0.5 M KOH, and then rinsed in deionized water (ddH2O). Then, it was dried and exposed to air plasma (Plasma Etch) for 3 minutes. They were then silanized by soaking in 0.1% dichlorodimethylsilane in *n*-heptane, followed by sonication in *n*-heptane and then in ethanol. 7 μl flow channels were constructed using two pieces of silanized cover glasses held together with double-sided tape and mounted into custom-machined coverslip holders ^47^. Channels were incubated with anti-TAMRA antibodies (diluted 1:100, Invitrogen) for 5 minutes, blocked with 5% Pluronic F-127 for 10 minutes, and then washed three times with BRB80 before incubating with microtubules. The temperature of the channel was set at 35°C using both an environmental chamber and an objective heater.

### Dynamic microtubule growth assay

We reconstituted microtubule growth from GMPCPP microtubule seeds to observe microtubule dynamics *in vitro* ^47^. GMPCPP microtubule seeds were prepared by polymerizing a 1:4 molar ratio of TAMRA-labelled to unlabelled tubulin in the presence of 1 mM of guanosine-5’-[(α,β)- methyleno]triphosphate (GMPCPP, Jena Biosciences, Germany), a non-hydrolyzable GTP analogue, in two cycles. Each day of the experiments, tubes of unlabelled and CF642R-labelled tubulin were thawed, mixed at a 1:17 molar ratio, aliquoted, and stored in liquid nitrogen. For consistency, one aliquot of this tubulin was used per experiment. To initiate an experiment, GMPCPP microtubule seeds were introduced into a channel and washed with BRB80.

Microtubule growth was then started by incubating flow channels with tubulin (final concentration 8 µM) in imaging buffer containing 1 mM GTP, 40 mM D-glucose, 64 nM catalase, 250 nM glucose oxidase, 10 mM dithiothreitol, 0.1 mg/mL bovine serum albumin (BSA) in BRB80, along with varying concentrations of the disordered domain (0, 5, 50, or 250 nM). The disordered domain was prepared fresh for every experiment. Low-retention tips and tubes were used throughout all experiments.

### Preparation of paclitaxel-stabilized GDP microtubules

Paclitaxel-stabilized GDP microtubules were prepared by polymerizing a 1:15 molar ratio of TAMRA-labelled: unlabelled tubulin with 4 mM MgCl_2_, 1 mM GTP, 5% DMSO in BRB80. The polymerization mixture was mixed and incubated at 37°C for 30 min. The microtubules were diluted with prewarmed BRB80 supplemented with 10 μM paclitaxel, pelleted at 100,000 rpm, 30 psi for 7 min using a Beckman-Coulter Airfuge, and resuspended in a suitable volume of BRB80 with 10 μM paclitaxel.

### Binding to paclitaxel-stabilized GDP and GMPCPP microtubules

A mixture of TAMRA-labelled paclitaxel-stabilized and GMPCPP double-cycled microtubules was introduced into flow channels and adhered to anti-TAMRA antibodies as described above. The paclitaxel-stabilized GDP microtubules had a lower ratio of TAMRA-labelled to unlabelled tubulin, resulting in dim microtubules compared to bright GMPCPP microtubules. All protein dilutions were prepared in BRB80 with 10 μM paclitaxel using low retention tips and tubes.

### Microtubule depolymerization assay

GMPCPP single-stabilized microtubules were polymerized using 25% labelled TAMRA tubulin in the presence of 1mM MgCl_2_, BRB80, and GMPCPP. The polymerization mixture was incubated on ice for 5 min, followed by incubation at 37°C for 2 hours. The microtubules were diluted with room-temperature BRB80, pelleted at 100,000 rpm and 30 psi for 5 min using a Beckman-Coulter Airfuge, and then resuspended in a suitable volume of warm BRB80. Protein concentrations of 5, 25, 50, and 500 nM of full-length CKAP2 and the disordered domain in a BRB80 imaging buffer without GTP. The prepared channels were sealed with nail polish gel to prevent drying. GMPCPP microtubules were imaged every 5 minutes during depolymerization for 2 hours and 30 minutes.

### Cell Culture

hTERT RPE-1 immortalized retinal pigment epithelial cells (ATCC number: CRL 4000) were a gift from Arnold Hayer (McGill University). Cells were cultured in DMEM/F-12 with high glucose and sodium pyruvate supplemented with 10% fetal bovine serum, 1% penicillin/streptomycin, and 10 µg/ml of hygromycin and incubated at 5% CO2 and 37°C.

### Cellular expression of CKAP2 and immunofluorescence

RPE-1 cells were cultured on either 8- (Ibidi µ-slide 8 well, 80807) or 18-well glass-bottom (Ibidi, µ-slide 18 well, 81817) plates prior to transfection. Transfections were performed using Xfect Transfection Reagent (Takara Bio, 631317) using 200-250 ng of DNA following the manufacturer’s instructions. Cells were fixed 24 h post-transfection with 100% methanol at - 20°C for 20 min. Cells were then blocked with 3% Bovine Serum Albumin (BSA) in 1x PBS overnight at 4 °C. Primary antibody incubation (rabbit polyclonal anti-TUBB3 - BioLegend, 802001, 1:200) was performed overnight at 4 °C. Cells were then washed three times with blocking solution and incubated with secondary antibody (AlexaFluor anti-rabbit 568, Invitrogen, A110011, 1:1000) for 1h30 at room temperature. Next, cells were washed three times with blocking solution and incubated with DAPI (1 µg/mL) for 15 minutes at room temperature. Microscopy was performed using a spinning disk microscope on a Quorum Diskovery platform, which was installed on a Leica DMi8 inverted microscope. This system consists of an HCX PL APO 63×/1.4 NA oil objective, a DISKOVERY multimodal imaging system (Spectral) with a multipoint confocal 50-µm pinhole spinning disk and dual iXon Ultra 512 × 512 EMCCD (Andor) cameras for simultaneous imaging, an ASI three-axis motorized stage controller, and an MCL Nano-view piezo stage, 488 nm, 561 nm, and 647 nm solid-state OPSL lasers linked to a Borealis beam conditioning unit. Image acquisition and microscope control were executed using MetaMorph (Molecular Devices).

### Image and data analysis

All images were processed and analyzed using Fiji (ImageJ) ^48^. Statistical analyses and charts were performed using GraphPad Prism 10.2.1 (GraphPad Software, Boston, Massachusetts, USA). In violin plots, the center line represents the median, the dashed lines represent upper and lower quartiles, and the dots represent independent measurements.

For numerical data of cell analysis, Shapiro–Wilk normality tests were applied, and either unpaired two-tailed t-tests or unpaired two-tailed Mann–Whitney U tests were applied. For categorical data in contingency tables, Fisher’s exact test was applied. Statistical significance was considered when *P* < 0.05.

To analyze the dynamic microtubule growth and depolymerization assays, the images were drift-corrected for stage drift using the TrackMate plugin ^49^, followed by a drift correction script (Hadim). Then, manually, the straight lines with a line width of 10 were drawn to cover microtubules. Using a kymograph builder plugin, all kymographs were generated from the lines and then analyzed. For the dynamic microtubule growth assay, the kymographs represented the growth distance (x-axis) over time (y-axis). Growth and shrinkage rates were manually and individually analyzed by drawing the lines and measuring the slopes of the growth or shrinkage. Catastrophe frequency was measured by counting the total number of catastrophe events on the kymographs over the total number of all microtubule growth within a channel. For the depolymerization assay, the kymographs showed microtubule depolymerization velocity (x-axis) over time (y-axis). To measure depolymerization velocity, the slope was analyzed for each individual microtubule. All *in vitro* experiment details are summarized in Supplementary Table 2.

All figures were prepared using Adobe Illustrator (29.0.1, Adobe Inc.).

## Acknowledgments

We thank the McGill Advanced Bioimaging Facility (ABIF) for training and technical support with microscopy experiments and Kim Munro (CRSB, McGill University) for assistance and analysis of circular dichroism (CD) spectroscopy experiments. Mass photometry (MP) work was conducted at the Structural Biology Platform of the Université de Montréal with the help of Normand Cyr, funded by a grant from the Canada Foundation for Innovation (#30574) and now supported by the Institut Courtois d’innovation biomédicale of the Université de Montréal. This work was supported by the Canadian Institutes of Health Research (CIHR) CIHR PJT-189995 and the Natural Sciences and Engineering Research Council of Canada (NSERC RGPIN-2024-05603) S.B. is a member of the Centre de recherche en biologie structurale, funded by Fonds de Recherche du Québec (Health Sector) Research Centres Grant #288558.

## Author contributions

TL, LP and SB conceptualized the study. TL, LP and SB provided the methodology. TL, LP and SB undertook the investigation. TL, LP and SB performed the visualization. SB acquired funding. SB supervised the study. SB and TL wrote the original draft of the manuscript. TL, LP and SB reviewed and edited the manuscript.

## Ethics declarations

### Data and materials availability

All data are available in the main text or the supplementary materials. Reagents will be provided upon request to the corresponding author.

## References

1. Wright, P. E. & Dyson, H. J. Intrinsically disordered proteins in cellular signalling and regulation. Nature Reviews Molecular Cell Biology 16, 18 (2015).

2. Guharoy, M., Pauwels, K. & Tompa, P. SnapShot: Intrinsic Structural Disorder. Cell 161, 1230–1230.e1 (2015).

3. Ward, J. J., Sodhi, J. S., McGuffin, L. J., Buxton, B. F. & Jones, D. T. Prediction and Functional Analysis of Native Disorder in Proteins from the Three Kingdoms of Life. Journal of Molecular Biology 337, 635–645 (2004).

4. Fonin, A. V. et al. Biological soft matter: intrinsically disordered proteins in liquid-liquid phase separation and biomolecular condensates. Essays Biochem 66, 831–847 (2022).

5. Wang, J. et al. A Molecular Grammar Governing the Driving Forces for Phase Separation of Prion-like RNA Binding Proteins. Cell 174, 688–699.e16 (2018).

6. Podolski, M., Mahamdeh, M. & Howard, J. Stu2, the Budding Yeast XMAP215/Dis1 Homolog, Promotes Assembly of Yeast Microtubules by Increasing Growth Rate and Decreasing Catastrophe Frequency *. Journal of Biological Chemistry 289, 28087–28093 (2014).

7. Alexandrova, V. V. et al. Theory of tip structure–dependent microtubule catastrophes and damage-induced microtubule rescues. Proceedings of the National Academy of Sciences 119, e2208294119 (2022).

8. Slautterback, D. B. CYTOPLASMIC MICROTUBULES : I. Hydra. Journal of Cell Biology 18, 367–388 (1963).

9. Mitchison, T. & Kirschner, M. Dynamic instability of microtubule growth. Nature 312, 237– 242 (1984).

10. Kirschner, M. & Mitchison, T. Beyond self-assembly: From microtubules to morphogenesis. Cell 45, 329–342 (1986).

11. Stepanova, T. et al. Visualization of Microtubule Growth in Cultured Neurons via the Use of EB3-GFP (End-Binding Protein 3-Green Fluorescent Protein). J. Neurosci. 23, 2655–2664 (2003).

12. Gomes Paim, L. M. & and Bechstedt, S. Regulation of microtubule growth rates and their impact on chromosomal instability. Cell Cycle 0, 1–20.

13. Salmon, E. D., Saxton, W. M., Leslie, R. J., Karow, M. L. & McIntosh, J. R. Diffusion coefficient of fluorescein-labeled tubulin in the cytoplasm of embryonic cells of a sea urchin: video image analysis of fluorescence redistribution after photobleaching. Journal of Cell Biology 99, 2157–2164 (1984).

14. Akhmanova, A. & Steinmetz, M. O. Control of microtubule organization and dynamics: two ends in the limelight. Nat Rev Mol Cell Biol 16, 711–726 (2015).

15. Howard, J. & Hyman, A. A. Microtubule polymerases and depolymerases. Curr Opin Cell Biol 19, 31–5 (2007).

16. Gangadharan, B., Kober, D. L. & Rice, L. M. A biochemical mechanism for Stu2/XMAP215-family microtubule polymerases. 2025.06.09.658552 Preprint at 10.1101/2025.06.09.658552 (2025).

17. Brouhard, G. J. et al. XMAP215 is a processive microtubule polymerase. Cell 132, 79–88 (2008).

18. Ayaz, P. et al. A tethered delivery mechanism explains the catalytic action of a microtubule polymerase. eLife Sciences 3, e03069 (2014).

19. Roostalu, J., Cade, N. I. & Surrey, T. Complementary activities of TPX2 and chTOG constitute an efficient importin-regulated microtubule nucleation module. Nature Cell Biology 17, 1422–1434 (2015).

20. Paim, L. M. G., Lopez-Jauregui, A. A., McAlear, T. S. & Bechstedt, S. The spindle protein CKAP2 regulates microtubule dynamics and ensures faithful chromosome segregation. Proceedings of the National Academy of Sciences 121, e2318782121 (2024).

21. McAlear, T. S. & Bechstedt, S. The mitotic spindle protein CKAP2 potently increases formation and stability of microtubules. eLife 11, e72202 (2022).

22. Emenecker, R. J., Griffith, D. & Holehouse, A. S. Metapredict: a fast, accurate, and easy-to-use predictor of consensus disorder and structure. Biophys J 120, 4312–4319 (2021).

23. Abramson, J. et al. Accurate structure prediction of biomolecular interactions with AlphaFold 3. Nature 630, 493–500 (2024).

24. Soltermann, F. et al. Quantifying Protein–Protein Interactions by Molecular Counting with Mass Photometry. Angewandte Chemie International Edition 59, 10774–10779 (2020).

25. Borgia, A. et al. Extreme disorder in an ultrahigh-affinity protein complex. Nature 555, 61– 66 (2018).

26. Amos, L. A. & Schlieper, D. Microtubules and Maps. in Advances in Protein Chemistry (ed. John M. Squire, and D. A. D. P.) vol. Volume 71 257–298 (Academic Press, 2005).

27. Kozlowski, L. P. Proteome-pI: proteome isoelectric point database. Nucleic Acids Research 45, D1112–D1116 (2017).

28. Arai, M., Suetaka, S. & Ooka, K. Dynamics and interactions of intrinsically disordered proteins. Current Opinion in Structural Biology 84, 102734 (2024).

29. Mollica, L. et al. Binding Mechanisms of Intrinsically Disordered Proteins: Theory, Simulation, and Experiment. Front. Mol. Biosci. 3, (2016).

30. Alderson, T. R., Pritišanac, I., Kolarić, Đ., Moses, A. M. & Forman-Kay, J. D. Systematic identification of conditionally folded intrinsically disordered regions by AlphaFold2. Proc Natl Acad Sci U S A 120, e2304302120 (2023).

31. Omidi, A., Møller, M. H., Malhis, N., Bui, J. M. & Gsponer, J. AlphaFold-Multimer accurately captures interactions and dynamics of intrinsically disordered protein regions. Proc Natl Acad Sci U S A 121, e2406407121 (2024).

32. Alushin, G. M. et al. High-Resolution Microtubule Structures Reveal the Structural Transitions in αβ-Tubulin upon GTP Hydrolysis. Cell 157, 1117–1129 (2014).

33. Mondal, S., Bonventre, E., Hancock, W. O. & Rice, L. M. Mechanisms of microtubule dynamics from single-molecule measurements. 2025.06.25.661545 Preprint at 10.1101/2025.06.25.661545 (2025).

34. Davey, N. E., Simonetti, L. & Ivarsson, Y. The next wave of interactomics: Mapping the SLiM-based interactions of the intrinsically disordered proteome. Curr Opin Struct Biol 80, 102593 (2023).

35. Puntervoll, P. et al. ELM server: a new resource for investigating short functional sites in modular eukaryotic proteins. Nucleic Acids Research 31, 3625–3630 (2003).

36. Davey, N. E. et al. Attributes of short linear motifs. Mol. BioSyst. 8, 268–281 (2011).

37. Holehouse, A. S. & Kragelund, B. B. The molecular basis for cellular function of intrinsically disordered protein regions. Nat Rev Mol Cell Biol 25, 187–211 (2024).

38. Van Roey, K. et al. Short Linear Motifs: Ubiquitous and Functionally Diverse Protein Interaction Modules Directing Cell Regulation. Chem. Rev. 114, 6733–6778 (2014).

39. Hornbeck, P. V. et al. PhosphoSitePlus, 2014: mutations, PTMs and recalibrations. Nucleic Acids Res 43, D512–520 (2015).

40. Cho, N. H. et al. OpenCell: Endogenous tagging for the cartography of human cellular organization. Science 375, eabi6983 (2022).

41. Meng, E. C. et al. UCSF ChimeraX: Tools for structure building and analysis. Protein Science 32, e4792 (2023).

42. Norholm, M. H. A mutant Pfu DNA polymerase designed for advanced uracil-excision DNA engineering. BMC Biotechnol 10, 21 (2010).

43. Bechstedt, S. & Brouhard, G. J. Doublecortin recognizes the 13-protofilament microtubule cooperatively and tracks microtubule ends. Dev Cell 23, 181–92 (2012).

44. Bitinaite, J. et al. USER friendly DNA engineering and cloning method by uracil excision. Nucleic Acids Res 35, 1992–2002 (2007).

45. Ashford, A. J., Andersen, S. L. A. & Hyman, A. A. Preparation of Tubulin from Bovine Brain. in Cell Biology, a Laboratory Handbook (ed. Celis, J. E.) vol. 2 205–212 (Academic Press, New York, 1998).

46. Hyman, A. et al. Preparation of modified tubulins. Methods Enzymol 196, 478–85 (1991).

47. Gell, C. et al. Microtubule dynamics reconstituted in vitro and imaged by single-molecule fluorescence microscopy. Methods Cell Biol 95, 221–45 (2010).

48. Schindelin, J., et al. Fiji: an open-source platform for biological-image analysis. Nat Methods 9, 676–682 (2012).

49. Tinevez, J.-Y. et al. TrackMate: An open and extensible platform for single-particle tracking. Methods 115, 80–90 (2017).

50. Pfleger, C. M. & Kirschner, M. W. The KEN box: an APC recognition signal distinct from the D box targeted by Cdh1. Genes Dev 14, 655–665 (2000).

