## Supplemental Material for "Enzymatic Function of an Intrinsically Disordered Protein"

**Extended Data Fig.1: Validation of the other three CKAP2 domains for microtubule.**

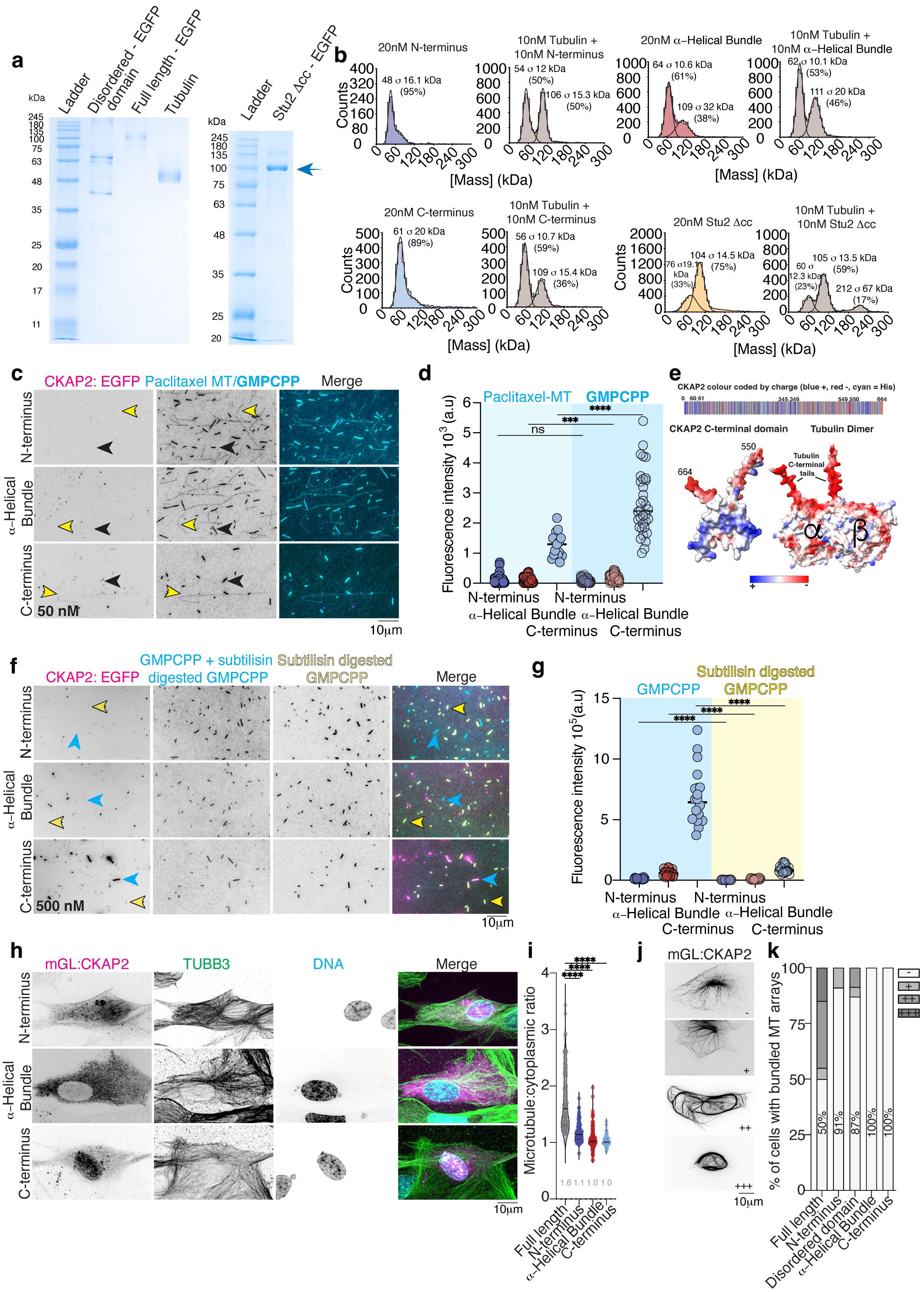

**a,** SDS-PAGE gel of 0.5 μg of purified tubulin, purified recombinant Disordered domain and full-length CKAP2, 1 μg of purified recombinant Stu2. **b,** Mass Photometry data for tubulin, CKAP2 domains (N-terminus, α-helical domain, C-terminus), and Stu2, both separately and together. **c,** Representative images for CKAP2 domains binding to paclitaxel and GMPCPP-stabilized microtubules at 50 nM. Yellow arrows highlight long, dim paclitaxel microtubules; black arrows indicate short, bright GMPCPP microtubules. (**d)** Quantification for data in (c). The black bar represents the mean; n varies from 17 to 50. *P* values are ns = 0.8719, *** = 0.0007, **** < 0.0001 (Mann-Whitney test, Welch's test was applied for analysis of C-terminus). **e,** CKAP2 linear protein schematic, colour-coded by charge (red- negative, blue- positive, cyan- Histidine) and AlphaFold3 model of the C-terminal domain showing charge clustering. **(f)** Representative images for binding of CKAP2 domains to control and subtilisin-digested microtubules at 500 nM. Yellow arrows show only subtilisin-digested microtubules; blue arrows indicate both control and subtilisin-digested microtubules. (**g**) Quantification for data in (f). The black bar represents the median, n = 20/construct. *P* values are **** < 0.0001 (Welch's test). **(h)** Representative images for three CKAP2 domains (N-terminus, α-helical domain, C-terminus) to microtubules in cells. **i,** Quantification for microtubule binding in cells. Full length n = 110 microtubules from 22 cells, N-terminus n = 115 microtubules from 23 cells. **** *P* < 0.0001, α-helical bundle N = 220 microtubules from 30 cells **** *P* < 0.0001, C-terminus n = 70 microtubules from 14 cells **** *P* < 0.0001. Mann-Whitney test. Values depicted in the chart refer to the median. **(j)** Representative images and rating for microtubule stabilization in cells (-) = no stabilized bundles, (+/++/+++) progressive amounts of stabilized bundles. **(k)** Quantification of cells with microtubule stability scores from (j) and Figure.1d. Full length n =

22 cells, N-terminus n = 23 cells *** $P = 0.0008$, disordered n = 23 cells * $P = 0.0225$, α-helical bundle n = 30 cells **** $P < 0.0001$, C-terminus n = 14 cells ** $P = 0.0058$. Fisher's exact test.

**Extended Data Fig.2: Shrinkage rates in the presence of the Disordered domain.**

**a**

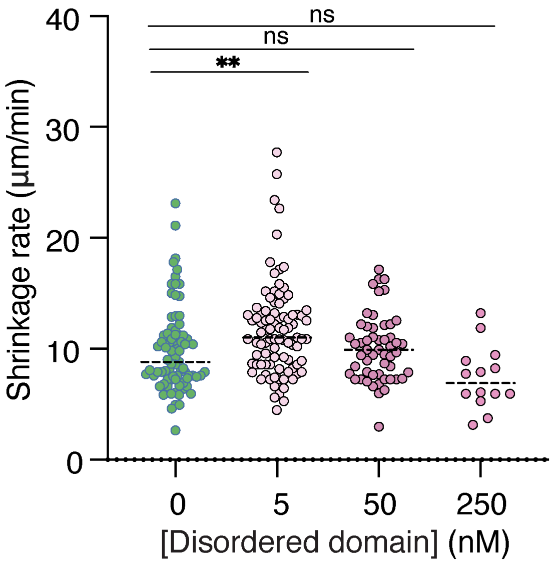


**a,** Quantification of microtubule shrinkage rates for data in Fig. 2c. (n = 78, 91, 53, 14 shrinkage events for 0, 5, 50 and 250 nM, respectively). The dashed black bar indicates the median (8.8, 11, 9.9, 6.9 for 0, 5, 50 and 250 nM) from at least 2 independent replicates. *P* values are indicated as ** = 0.0038, ns > 0.9999, ns = 0.172 (Kruskal-Wallis test).

**Extended Data Fig.3: Sequence conservation analysis of CKAP2.**

**a**

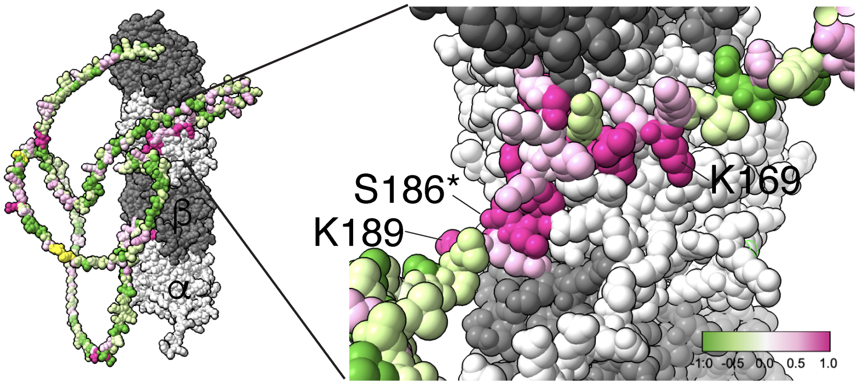


**b**

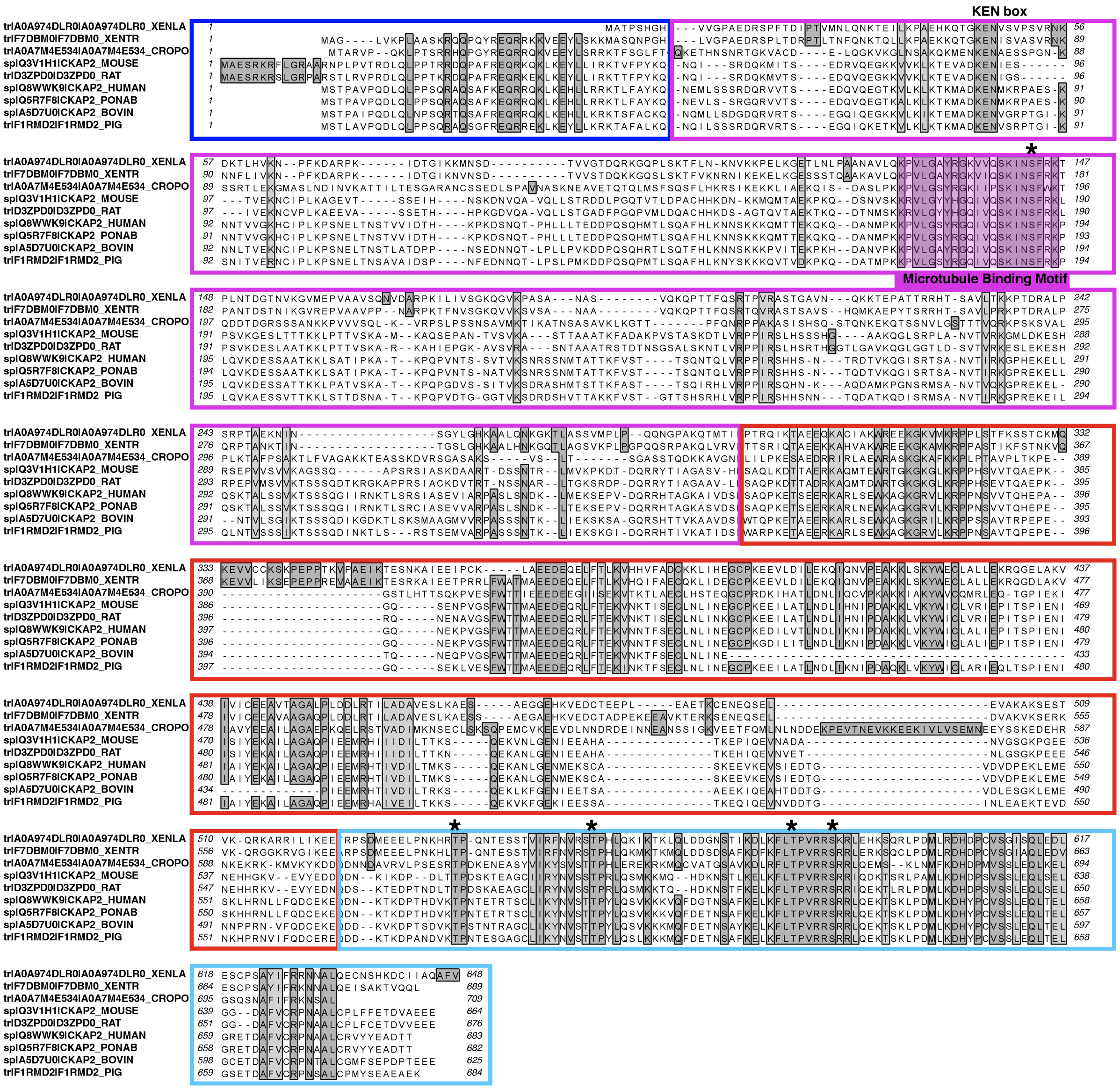

**a,** AlphaFold3 model of the mouse CKAP2, colour-coded by conservation (magenta conserved, green variable, yellow unique for mouse CKAP2) and enlargement of the microtubule binding segment. **b,** Sequence alignment for CKAP2 from Xenopus laevis, Xenopus tropicalis, Saltwater crocodile, mouse, rat, human, orangutan, bovine and pig CKAP2. Domains are colour-coded. The microtubule-binding domain, KEN box, a signal for ubiquitination and degradation by the APC [50], and conserved phosphorylation sites (*) are labelled.

**Supplementary Table 1: The CKAP2 domains were amplified using the following primers**

| Name of a fragment | Forward primer, 5' → 3' | Reverse primer, 5' → 3' |
|---|---|---|
| **The N-terminal α-helix** | GGGGAGCUATGGCAGAGTCCAGGAAACG | GGGGAACUATTTTCTTGTTTGTATGGAAACACTG |
| **The Disordered domain** | GGGGAGCUCAGATATCCAGAGATCAGAAAATG | GGGGAACUCTCTATGAACACTTGCTCCTGC |
| **The central α-helical bundle** | GGGGAACUTTCCTCTTCCGCTACGTCGG | GGGGAACUTTGATCATCTTCATACACCTCAAC |
| **The C-terminal** | GGGGAACUTTCCTCTTCCGCTACGTCGG | GGGGAACUTTGATCATCTTCATACACCTCAAC |

### Supplementary Table 2: Summary of *in vitro* experiments

| Name of an assay | Name of the fragment | Concentration, nM | n = microtubules | | Mean | | Statistics |
|---|---|---|---|---|---|---|---|
| ***in vitro* microtubule binding (Fig. 1H and fig. S1D)** | | | *GMPCPP* | *Paclitaxel* | *GMPCPP* | *Paclitaxel* | |
| | Full length | 5 | 50 | 32 | $0.71 \bullet 10^{5}$ | $0.99 \bullet 10^{5}$ | |
| | | 25 | 50 | 46 | $0.51 \bullet 10^{6}$ | $0.68 \bullet 10^{6}$ | |
| | | 50 | 50 | 45 | $0.85 \bullet 10^{6}$ | $0.94 \bullet 10^{6}$ | |
| | | 125 | 50 | 46 | $1.14 \bullet 10^{6}$ | $1.45 \bullet 10^{6}$ | |
| | | 250 | 50 | 41 | $1.47 \bullet 10^{6}$ | $1.54 \bullet 10^{6}$ | |
| | | 500 | 50 | 43 | $1.55 \bullet 10^{6}$ | $1.96 \bullet 10^{6}$ | |
| | Disordered Region | 5 | 47 | 44 | $0.37 \bullet 10^{5}$ | $0.13 \bullet 10^{5}$ | |
| | | 25 | 50 | 50 | $1.35 \bullet 10^{5}$ | $0.70 \bullet 10^{5}$ | |
| | | 50 | 50 | 50 | $3.22 \bullet 10^{5}$ | $2.73 \bullet 10^{5}$ | |
| | | 125 | 48 | 50 | $4.52 \bullet 10^{5}$ | $4.25 \bullet 10^{5}$ | |
| | | 250 | 43 | 50 | $5.04 \bullet 10^{5}$ | $4.99 \bullet 10^{5}$ | |
| | | 500 | 47 | 48 | $5.82 \bullet 10^{5}$ | $5.92 \bullet 10^{5}$ | |
| | | | *GMPCPP* | *Paclitaxel* | *GMPCPP* | *Paclitaxel* | Mann-Whitney test (Welch's t test for C – terminus) |
| | N-terminus | 50 | 50 | 50 | 86.0 | 105.80 | ns 0.8719 |
| | Helical Bundle | 50 | 50 | 50 | 170.50 | 101.67 | *** 0.0007 |
| | C-terminus | 50 | 34 | 17 | 2651.89 | 1261.67 | ****< 0.0001 |
| **GMPCPP and Subtilisin digested GMPCPP (fig. S1G)** | | | *GMPCPP* | *SubtilisinD igested GMPCPP* | *GMPCPP* | *Subtilisin Digested GMPCPP* | Welch's t test |
| | N-terminus | 500 | 21 | 20 | $1.21 \bullet 10^{4}$ | $0.34 \bullet 10^{4}$ | ****< 0.0001 |
| | Helical Bundle | 500 | 20 | 21 | $0.59 \bullet 10^{5}$ | $0.11 \bullet 10^{5}$ | ****< 0.0001 |
| | C-terminus | 500 | 20 | 22 | $6.79 \bullet 10^{5}$ | $0.91 \bullet 10^{5}$ | ****< 0.0001 |
| **Dynamic Assay Microtubule growth (Fig. 2D)** | Control | 0 | 97 | | 0.47 | | Kruskal-Wallis test |
| | Disordered Region | 5 | 102 | | 0.57 | | *** 0.0003 |
| | | 50 | 78 | | 0.67 | | ****< 0.0001 |
| | | 250 | 38 | | 0.98 | | ****< 0.0001 |
| **Dynamic Assay Shrinkage Rate (fig. S2A)** | Control | 0 | 78 | | 8.8 | | Kruskal-Wallis test |
| | Disordered Region | 5 | 91 | | 11 | | ** 0.0038 |
| | | 50 | 53 | | 9.9 | | ns > 0.9999 |
| | | 250 | 14 | | 6.9 | | ns 0.1729 |
| **Dynamic Assay Catastrophe Frequency (Fig. 2E)** | Control | 0 | 78 | | 0.040 | | |
| | Disordered Region | 5 | 91 | | 0.067 | | |
| | | 50 | 53 | | 0.029 | | |
| | | 250 | 14 | | 0.018 | | |
| **Depolymerization assay (Fig. 2H)** | Control | 0 | 56 | | 0.41 | | Kruskal-Wallis test |
| | Full length | 5 | 59 | | 0.81 | | **** < 0.0001 |
| | | 25 | 47 | | 0.59 | | * 0.0117 |
| | | 50 | 22 | | 0.28 | | ns 0.2035 |
| | | 500 | 59 | | 0.18 | | **** < 0.0001 |
| | Control | 0 | 46 | | 0.39 | | Kruskal-Wallis test |
| | Disordered Region | 5 | 54 | | 0.98 | | ****< 0.0001 |
| | | 25 | 36 | | 0.58 | | ** 0.0039 |
| | | 50 | 79 | | 0.42 | | ns > 0.9999 |
| | | 500 | 24 | | 0.19 | | ** 0.0030 |

Data for graphs were plotted from at least two independent replicates.
